# Direct monitoring of cAMP at the cardiac ryanodine receptor using a novel targeted fluorescence biosensor mouse

**DOI:** 10.1101/623934

**Authors:** Filip Berisha, Konrad R. Götz, Jörg W. Wegener, Christiane Jungen, Ulrike Pape, Axel E. Kraft, Svenja Warnke, Diana Lindner, Dirk Westermann, Stefan Blankenberg, Christian Meyer, Gerd Hasenfuß, Stephan E. Lehnart, Viacheslav O. Nikolaev

## Abstract

**Rationale:** Cyclic adenosine monophosphate (cAMP) is a ubiquitous second messenger which, upon β-adrenergic receptor (β-AR) stimulation, acts in microdomains to regulate cardiac excitation-contraction coupling by activating the cAMP-dependent protein kinase (PKA) phosphorylation of calcium handling proteins. One crucial microdomain is in vicinity of the cardiac ryanodine receptor type 2 (RyR2) which is associated with arrhythmogenic diastolic calcium leak from the sarcoplasmic reticulum (SR) often occurring upon RyR2 hyperphosphorylation by PKA and calcium/calmodulin-dependent kinase.

**Objective:** We sought to establish a real time approach capable of directly visualizing cAMP and its pathological changes in the vicinity of RyR2 by generating a proper targeted biosensor and transgenic mouse model to express it in adult cardiomyocytes.

**Methods and Results:** We generated transgenic mice expressing a novel targeted fluorescent biosensor for RyR2-associated cAMP in adult mouse cardiomyocytes. In healthy cardiomyocytes, β_1_-AR but not β_2_-AR stimulation strongly increased local RyR2-associated cAMP levels. However, in cardiac hypertrophy induced by aortic banding, there was a marked subcellular redistribution of phosphodiesterases (PDEs) 2, 3 and 4, which included a dramatic loss of the local pool of PDE4. This was also accompanied by measurable β_2_-AR-induced cAMP signals, increased SR calcium leak and arrhythmia susceptibility.

**Conclusions:** Our new targeted biosensor expressed in transgenic mice can visualize cAMP levels in the vicinity of cardiac RyR2 in healthy and diseased cardiomyocytes. In the future, this novel biosensor can be used to better understand alterations of RyR2-associated cAMP in cardiovascular diseases and local actions of new therapies.

## Introduction

β-Adrenergic receptors (β-ARs) regulate cardiac excitation-contraction coupling by stimulation of the second messenger 3’,5’-cyclic adenosine monophosphate (cAMP) synthesis and downstream phosphorylation of several calcium handling proteins by the cAMP-dependent protein kinase (PKA). These include L-type calcium channels responsible for calcium influx upon depolarization in the T-tubular membrane invaginations, cardiac ryanodine receptors type 2 (RyR2s) responsible for calcium-induced calcium release from the sarcoplasmic reticulum (SR), and activation of the SR calcium ATPase (SERCA2a) via phospholamban phosphorylation.^1, 2^ Individually, each of these calcium handling proteins exists as a macromolecular complex comprised of A-kinase anchoring proteins, PKA, phosphatases, cAMP degrading enzymes phosphodiesterases (PDEs) and additional components, in concert enabling differentially regulated subcellular cAMP microdomains.^3–6^

Under physiological conditions, these microdomains tightly regulate calcium cycling and fine-tune the classical “fight-or-flight” response. In contrast, cardiac diseases such as heart failure lead to significant molecular remodeling of the signaling complexes and alterations in microdomain-specific cAMP dynamics.^4, 7^ For example, in the healthy heart, RyR2 channels are usually closed in diastole and stabilized by allosteric interactions with the small regulatory protein calstabin2 (also called FKBP12.6)^8^ and PDE4, which lowers cAMP in the vicinity of RyR2 and limits PKA-dependent phosphorylation in a negative feedback loop.^9^ In heart failure, however, calstabin2 and PDE4 are decreased in the RyR2 complex, leading to hyperphosphorylation of RyR2 and increased diastolic calcium leak, increasing the susceptibility for life-threatening arrhythmias.^9, 10^ In addition, RyR2 is phosphorylated by the calcium-calmodulin dependent protein kinase II (CaMKII) downstream of β_1_-AR subtype, which also increases calcium leak due to RyR2 hyperphosphorylation^11^ e.g. via the exchange protein directly activated by cAMP (Epac).^12, 13^ Although these disease-associated, chronic molecular changes have been well established,^14^ direct microdomain-specific monitoring of cAMP in the vicinity of RyR2 with high spatial and temporal resolution which could allow detection of early disease-relevant mechanisms, was so far not possible due to a lack of appropriate imaging techniques.

Here, we designed the first targeted cAMP biosensor, which can be used to directly monitor local cAMP dynamics in the RyR2 microdomain. Using this sensor in transgenic mice subjected to an *in vivo* hypertrophy model of early cardiac disease, we could observe dramatic pathology-associated changes in the microdomain-specific cAMP levels. Furthermore, there were functionally relevant changes in local cAMP degradation presumably due to subcellular redistribution of PDE2, 3 and 4 isoforms. Therefore, our novel targeted fluorescence biosensor and the respective transgenic mouse model should be useful for understanding real time cAMP dynamics at the cardiac RyR2 under normal and pathological conditions.

## Results

### Generation of a new RyR2-targeted biosensor transgenic mouse model

To monitor cAMP dynamics locally in the vicinity of RyR2, we sought to target the FRET-based cAMP biosensor Epac1-camps to the macromolecular complex of the calcium release unit without affecting cardiac function. Fusing the biosensor to the cytoplasmic N-terminus of junctin (JNC), a transmembrane SR protein which forms a stable complex with RyR2, resulted in a new targeted cAMP sensor termed Epac1-JNC. Next, we generated transgenic mice that express this new biosensor in adult myocardium under the control of α-myosin heavy chain promoter (**Fig. 1A**). Confocal microscopy of transgenic adult mouse ventricular cardiomyocytes immunostained with RyR2 antibody revealed proper co-localization of the biosensor with subcellular calcium release units (**Fig. 1B**). Importantly, this targeted biosensor could be co-immunoprecipitated with RyR2 (**Supplementary Fig. 1**), and live cell FRET imaging in isolated adult ventricular myocytes showed a rapid FRET response to the beta-adrenergic agonist isoproterenol (ISO), indicating cAMP synthesis and accumulation at RyR2 clusters (**Fig. 1C**).

**Figure 1.**
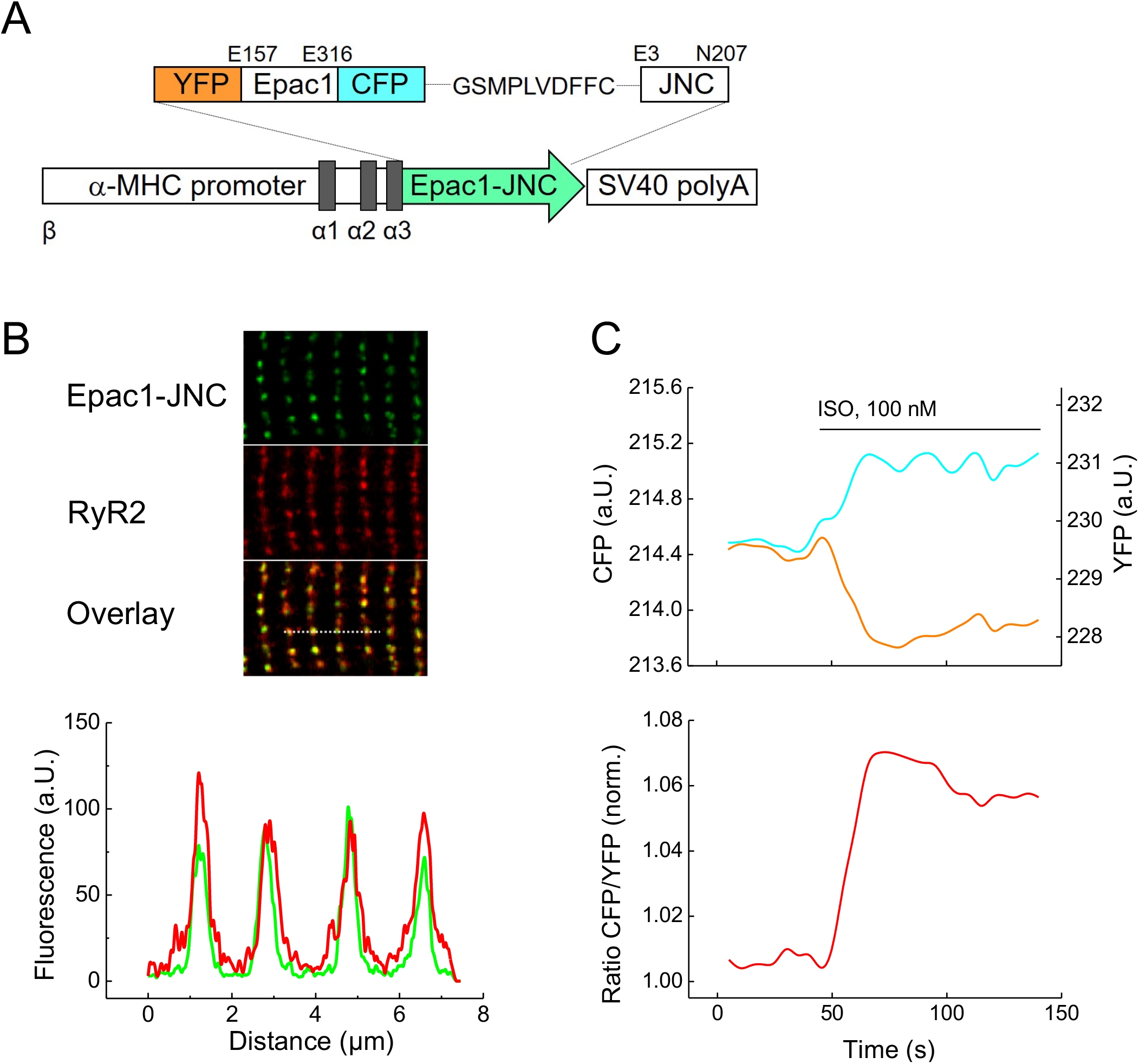
Generation of Epac1-JNC transgenic mice. (**A**) Schematic representation of the Epac1-JNC sensor construct which includes two fluorophores, yellow (YFP) and cyan fluorescent proteins (CFP), flanking the cAMP binding domain from Epac1 and fused to the N-terminus of junctin (JNC). Epac1-JNC was transgenically expressed in mice under the control of the cardiomyocyte specific α-MHC promoter. (**B**) Representative confocal image of a transgenic Epac1-JNC cardiomyocyte immunostained with RyR2 antibody. Epac1-JNC and RyR2 colocalization is confirmed by the intensity overlay of both fluorescent signals measured across the bar, 8 μm. (**C**) Representative single YFP and CFP intensities and CFP/YFP ratio trace (n=10) of Epac1-JNC cardiomyocyte upon stimulation with isoproterenol (ISO). An increase in the FRET ratio represents an increase in local cAMP.

Epac1-JNC mice had normal life expectancy and did not show any phenotypic abnormalities. The heart and left ventricular weights divided by body mass or tibia length, respectively, were comparable to wildtype littermates (**Supplementary Fig. 2A,B**) as were the morphology of cardiac cross-sections and cardiomyocyte size (**Supplementary Fig. 2C,D**). Echocardiography at 6 months of age showed no signs of ventricular dilation or impairment of contractile function. There were no significant differences in heart dimensions compared to wildtype littermates apart from a slight increase of diastolic but not systolic anterior and posterior wall thicknesses (**Supplementary Table 1**). Notably, this contrasts with the previously described pronounced hypertrophy phenotype in transgenic mice overexpressing canine junctin >10-30 fold.^15, 16^ In our model, based on qPCR, junctin overexpression was less than 30-fold (**Supplementary Fig. 2E**). Apparently, the mouse Epac1-JNC transgene does not lead to the dramatic cardiac phenotype typical for canine junctin transgenic lines.

### cAMP signals in the RyR2 microdomain after β_1_- and β_2_-adrenoceptor stimulation

To monitor local cAMP dynamics at RyR2 and to understand how it might be affected in early cardiac disease, we isolated ventricular cardiomyocytes from healthy (sham operated) mice and animals 8-10 weeks after transverse aortic constriction (TAC) used as a model of functionally compensated cardiac hypertrophy in the FVB/N1 background^4^ (**Supplementary Table 2**). Freshly isolated and plated cells were stimulated with a saturating concentration of the β-adrenergic agonist ISO (100 nmol/L) in combination with selective β_2_- or β_1_-AR blockers (ICI 118551 at 50 nmol/L or CGP 20712 A at 100 nmol/L, respectively)^17,18^ to enable β_1_-versus β_2_-specific receptor activation. Interestingly, while β_1_-AR stimulation produced comparable signals in sham and TAC cells, β_2_-AR activation showed no cAMP increase in the RyR2 microdomain under healthy conditions (**Fig. 2A-C**). This suggests that either the number of β_2_-ARs is insufficient in these cells^19^ or cAMP synthesis activated by β_2_-AR in T-tubular invaginations^20^ near RyR2 clusters does not cross the dyadic cleft to reach the RyR2 microdomain. In sharp contrast, TAC myocytes showed a clear increase of local cAMP directly in the RyR2 microdomain after β_2_-AR stimulation (**Fig. 2D**). In contrast to the lack of changes observed post-TAC for β_1_-AR stimulation, β_2_-AR stimulation induced a pronounced cAMP response in the RyR2 microdomain (**Fig. 2E**). To study the responsible molecular mechanisms, we next analyzed the effects of different PDE isoforms expressed in the heart.

**Figure 2.**
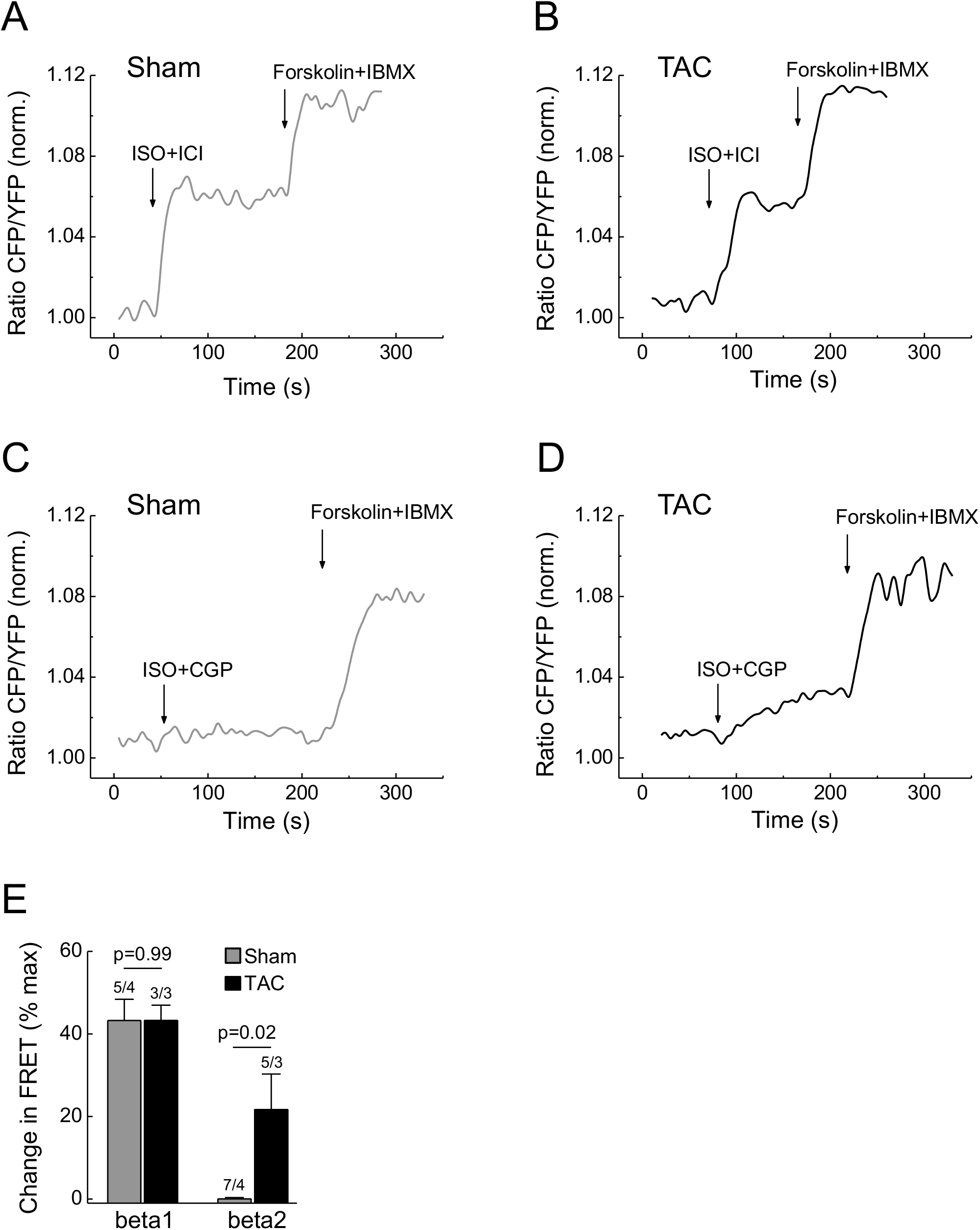
β_2_-adrenergic receptor stimulation increases local cAMP at RyR2 in diseased but not in healthy cells. (**A,B**) Representative FRET traces from sham and TAC Epac1-JNC cardiomyocytes selectively stimulated with β_1_-AR ligands (100 nmol/L ISO plus 50 nmol/L of the β_2_-AR blocker ICI 118551). (**C,D**) Representative traces from sham and TAC Epac1-JNC myocytes after selective β_2_-AR stimulation (100 nmol/L ISO plus 100 nmol/L of the β_1_-AR blocker CGP 20712A). β_2_-AR shows no effect on local cAMP in healthy but substantial increase in diseased cells. Subsequent maximal stimulation of adenylyl cyclase by 10 μmol/L of forskolin and inhibition of PDEs by 100 μmol/L IBMX was used to obtain the maximal FRET response. (E) Quantification of the FRET experiments shown in A-D. Means ± SE, number of cells/hearts is indicated above the bars. * - significant differences at p<0.05, mixed ANOVA followed by Wald χ^2^ test.

### Differential PDE regulation of the RyR2 microdomain

To understand how local cAMP levels are controlled in the RyR2 microdomain compared to the bulk cytosol, we isolated cardiomyocytes either from untreated Epac1-JNC or Epac1-camps transgenic mice^21^ and stimulated them with a subsaturating concentration of ISO (3 nmol/L) followed by selective inhibition of PDE2, 3 or 4 subfamilies. Interestingly, while PDE2 inhibition did not influence cAMP signaling in the RyR2 microdomain (**Fig. 3A,B**) and the PDE3 inhibitor effect was relatively small, the PDE4 inhibitor rolipram produced a very strong local signal increase in the RyR2 microdomain (**Fig. 3C**). Of note, the actual cAMP affinity of the Epac1-JNC sensor measured using a cell permeable cAMP analogue was comparable with that of the cytosolic sensor Epac1-camps (**Supplementary Fig. 3**). On average, the PDE4-dependent signal was 2-3-fold stronger in the RyR2 microdomain compared to the cytosol (**Fig. 3C-E**). This supports the concept that PDE4 anchored directly to the RyR2 complex represents the major cAMP-degrading activity involved in the local regulation of RyR2 microdomains.^9^

**Figure 3.**
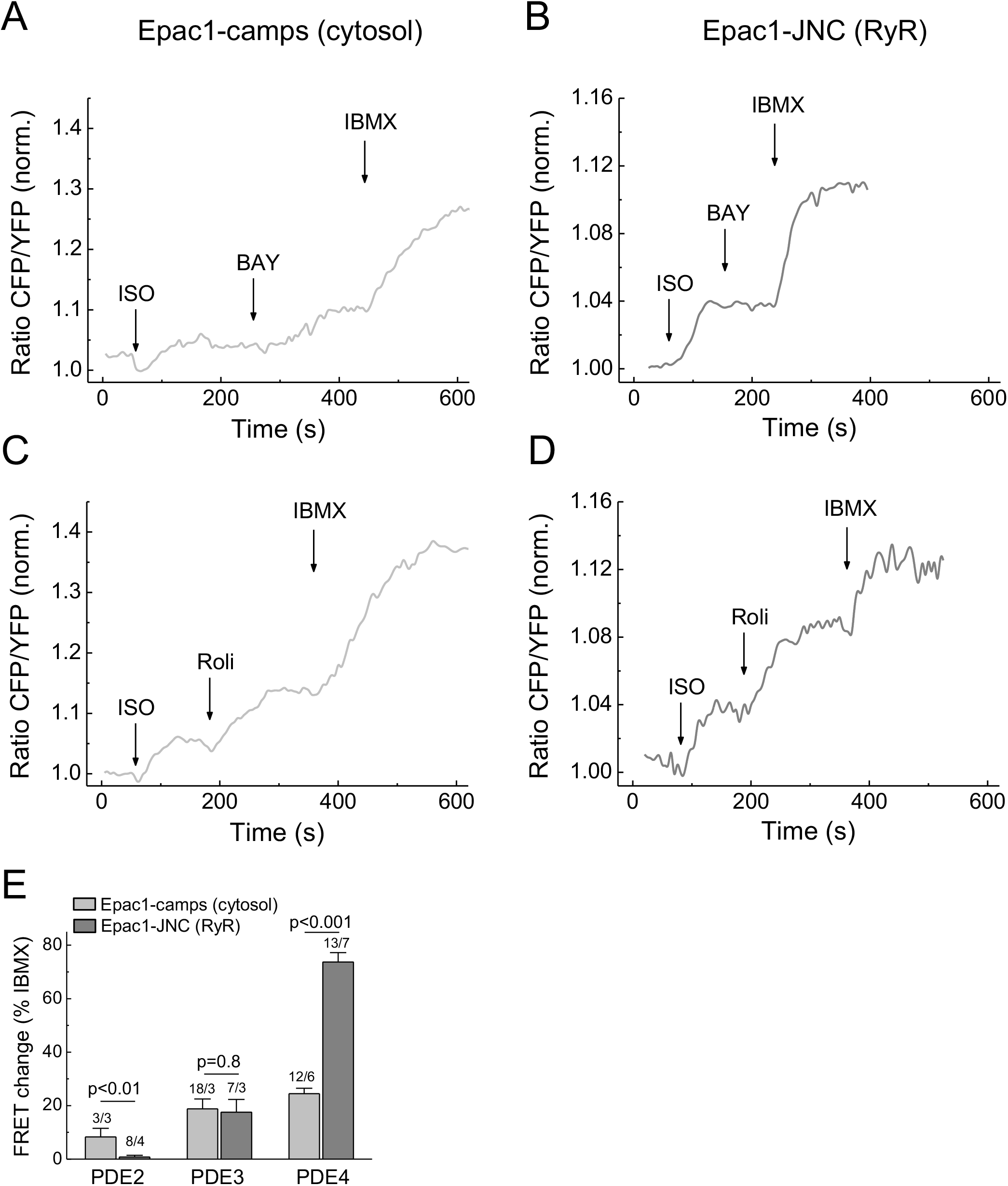
Contributions of various PDE families to local cAMP hydrolysis at the RyR2 after β-AR stimulation. Representative FRET traces from cardiomyocytes expressing Epac1-camps (**A,C**) or Epac1-JNC (**B,D**) treated with 3 nmol/L ISO for submaximal β-AR stimulation and subsequently either with the PDE2 inhibitor BAY 60-7550 (BAY, 100 nmol/L) (A,B) or with the PDE4 inhibitor rolipram (Roli, 10 μmol/L) (C,D). In the RyR2 microdomain, we could detect virtually no effect of the PDE2 inhibitor, while PDE4 contribution was significantly higher than in the bulk cytosol. (**E**) Quantification of the FRET experiments shown in A-D and similar experiments performed using the PDE3 inhibitor cilostamide (10 μmol/L). Means ± SE, number of hearts/cells is indicated above the bars. * - significant differences at p<0.05, mixed ANOVA followed by Wald χ^2^ test.

### Microdomain specific effects of PDEs are significantly changed in early heart disease

To analyze whether the local β-AR/cAMP signal changes observed after TAC might originate from PDE changes in the RyR2 microdomain, we performed FRET measurements comparing sham and TAC transgenic Epac1-JNC cells. Cardiac remodeling post-TAC led to a significant increase of PDE2, but a smaller increase of PDE3 inhibitor effect in the RyR2 microdomain (**Fig. 4A,B,E**). Strikingly, while the dominant PDE4 contribution was confirmed for sham cells, cAMP synthesis was dramatically blunted after TAC in the RyR2 microdomain (**Fig. 4C-E**). However, whole-cell expression levels of all three PDEs detected by immunoblotting were not altered at this early stage of disease (**Supplementary Fig. 4**). This strongly suggests that TAC leads to local alterations of PDE activities in the RyR2 microdomain, especially to a local depletion of PDE4, which were previously not detectable using classical biochemical techniques, somewhat similar to previously observed PDE redistribution in subsarcolemmal microdomains.^4^ To further support the notion that PDE4 may locally restrict microdomain access of β_2_-AR/cAMP to RyR2, we pre-blocked PDE4 selectively in healthy cells with 10 μM rolipram and stimulated β_2_-AR as described above (see **Fig 2C-E**). In this case, β_2_-AR stimulation produced a clear FRET response, which amounted to 12.8±1.2 % of the maximal response to forskolin plus IBMX (mean ± SE, n=12 cells from N=3 mice; compare to **Fig. 2E**).

**Figure 4.**
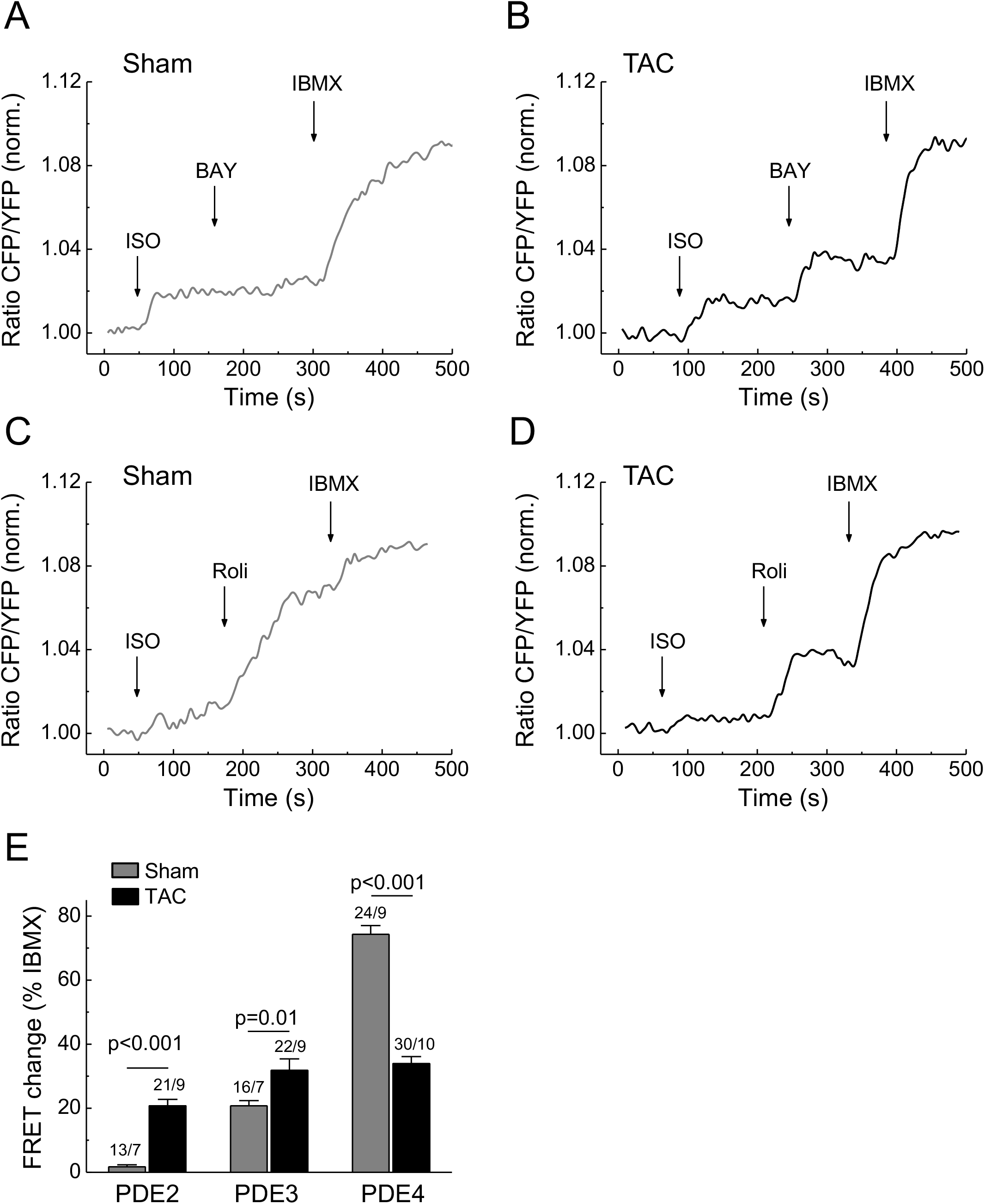
Contribution of PDEs to local cAMP control in diseased cardiomyocytes. Representative FRET traces from Epac1-JNC cells isolated from either sham (**A,C**) or TAC mice (**B,D**) and treated as described in Fig. 3. Diseased cells show an increase in PDE2 and PDE3 effects in the microdomains, while the predominant contribution of PDE4 is dramatically reduced. (**E**) Quantification of the FRET experiments shown in A-D and similar experiments performed with the PDE3 inhibitor cilostamide (10 μmol/L). Means ± SE, number of hearts/cells is indicated above the bars. * - significant differences at p<0.05, mixed ANOVA followed by Wald χ^2^ test.

### In hypertrophy, β_2_-adrenoceptor stimulation increases arrhythmia susceptibility

To answer the question whether the changes observed in hypertrophic cardiomyocytes with PDEs redistributed in RyR2 microdomains and augmented local β_2_-AR/cAMP signaling have an impact on calcium leak and arrhythmogenesis, we investigated the corresponding local calcium signals in cardiomyocytes. Confocal line-scan imaging immediately following electrical pacing of sham cells revealed that selective β_2_-AR versus non-selective β-AR stimulation had virtually no effect versus a significant increase in calcium spark generation (**Fig. 5B**). In contrast, post-TAC cardiomyocytes showed a significantly increased calcium spark frequency (**Fig. 5A,B**). Furthermore, in post-TAC but not in sham cells we observed a faster calcium transient decay upon β_2_-AR stimulation (**Supplementary Fig. 5A and 4B**). This was accompanied by faster cell shortening, as confirmed by β_1_-AR stimulation of sham cells (**Supplementary Fig. 5E**). These data strongly suggest that a PDE redistribution post-TAC augments β_2_-AR/cAMP levels in the RyR2 microdomain while SR calcium loading via SERCA2a is accelerated.

**Figure 5.**
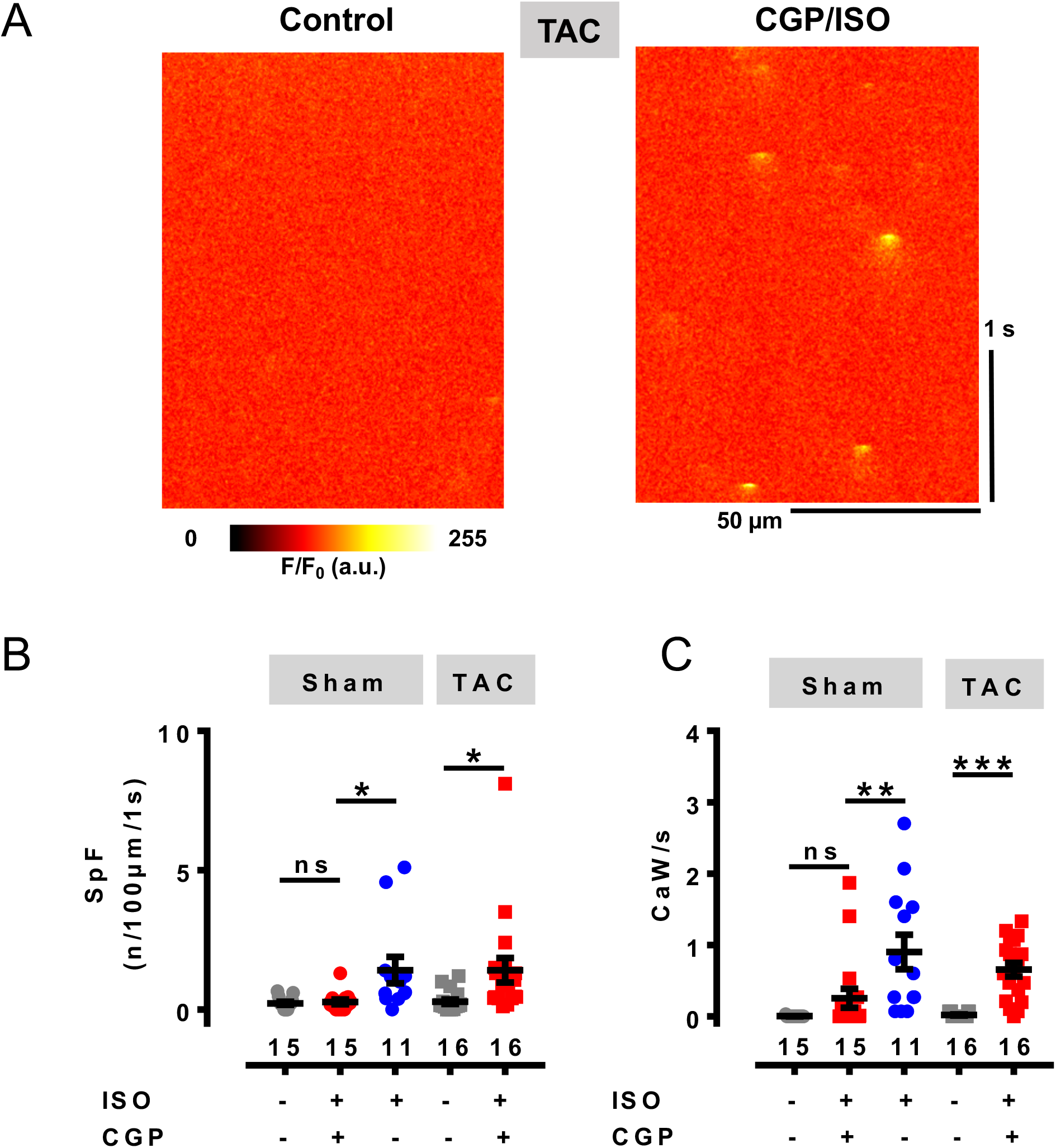
Analysis of Ca^2+^ sparks in cardiomyocytes from sham and TAC mice. (**A**) Original line scan confocal images of a cardiomyocyte from a TAC heart without and with ISO plus CGP 20712A (both 100 nmol/L) measured immediately after pacing (1 Hz). (**B**) Ca^2+^ spark frequency (SpF) analysis in cardiomyocytes from sham and TAC mice under control conditions, in the presence of ISO plus CGP 20712A (red), and in the presence of ISO (blue, sham only). Calcium sparks were analyzed after 5 consecutive Ca^2+^ transients elicited by field stimulation at 1Hz. (**C**) Ca^2+^ wave (CaW) frequency in cardiomyocytes from sham and TAC hearts under control conditions, in the presence of ISO plus CGP 20712A (red), and in the presence of ISO (blue, sham only). The number of CaWs per second following 1 Hz field stimulation is shown. Significant differences calculated by paired Student’s t-test are indicated: *, p<0.05; **, p<0.01; ***, p<0.001; ns, not significant. Graphs B and C show each data point and the mean ± SE. Numbers represent number of cardiomyocytes isolated from 3 mouse hearts for each group.

To assess if the increased calcium leak and faster calcium loading in post-TAC cardiomyocytes lead to increased arrhythmia susceptibility, we imaged calcium waves following β_2_-AR stimulation under steady-state conditions immediately following 1Hz pacing. While untreated post-TAC cardiomyocytes typically showed no calcium waves similar to sham, β_2_-AR stimulation induced a significant increase in calcium waves only in post-TAC but not in sham cells (**Fig. 5C**). Furthermore, ISO stimulation resulted in a profound increase in calcium waves in the same sham cells that showed no significant effect to β_2_-AR selective stimulation (**Fig. 5C**). Interestingly, in contrast to sham, β_2_-AR stimulation induced repetitive calcium wave events in most post-TAC cardiomyocytes, indicating that RyR2 dysfunction resulted in diastolic calcium overload, automaticity, and triggered arrhythmias.

To assess whether this translates into ventricular arrhythmias, we performed Langendorff experiments with electrically stimulated TAC and sham hearts perfused with ISO plus the β_1_-AR blocker CGP 20712A. Compared to sham, TAC hearts showed a very strong tendency towards increased number of premature ventricular complexes under these conditions (**Supplementary Fig. 6**).

## Discussion

In this work, we designed a novel microdomain-specific, targeted FRET biosensor and generated the first transgenic mouse model to measure real time cAMP dynamics in the direct vicinity of cardiac RyR2 channels. To target the highly sensitive cytosolic Epac1-camps sensor to this important microdomain, we sought to obtain a localized fusion protein, which forms a stable macromolecular complex with RyR2. Since the size of calcium release channel itself (megadalton range) makes a fusion protein strategy difficult, an essential RyR2 interacting protein may serve as targeting candidate. However, one possible protein with high-affinity interactions, calstabin2 (FKBP12.6), previously used to target a calcium sensor to the diadic cleft,^22^ was shown to dissociate from RyR2 upon β-adrenergic stimulation^23^ investigated here. Therefore, we used the mouse junctin homologue as interaction partner to target the RyR2 complex^24^ through a fusion protein strategy with Epac1-camps. The new RyR2-targeted biosensor Epac1-JNC (see **Fig. 1A**) showed excellent co-localization with RyR2 in adult cardiomyocytes and generated robust FRET responses, which were clearly different from those measured in the cytosol (see **Figs. 2 and 3**). This suggests that the novel targeted biosensor can specifically report local cAMP dynamics in the RyR2 microdomain. Importantly, biosensor affinity for cAMP was comparable with that of the parental cytosolic construct (**Supplementary Fig. 3**), indicating that local versus global cAMP responses can be roughly compared between the respective mouse strains.

Since overexpression of a canine JNC isoform in transgenic mice led to marked cardiac hypertrophy,^15, 16^ we thoroughly investigated whether Epac1-JNC mice might have any adverse phenotype. Up to the age of 6 months, there were no changes in cardiomyocyte size, cardiac morphometric or echocardiographic parameters, apart from a slight increase of posterior but not anterior diastolic wall thickness (see **Supplementary Fig. 2** and **Supplementary Table 1**). This small difference did not significantly affect development of cardiac hypertrophy after TAC in Epac1-JNC (see **Supplementary Table 2**), which was comparable to the phenotype observed in wildtype FVB/N1 mice (**Supplementary Table 3**), and also to that previously described in Epac1-camps mice which underwent the same procedure.^5^ Here, the transcript level of mouse JNC expression in Epac1-JNC mice was less than 30-fold times control, which is comparable with the previously described canine JNC transgenic models (**Supplementary Fig. 2E**).^15, 16^

Surprisingly, β_2_-AR, which was previously located in the T-tubules of rat ventricular myocytes where it generates highly compartmentalized second messenger signals^20^, did not produce any increase in local RyR2-associated cAMP levels in healthy mouse cells (see **Fig. 2**). This suggests that either the expression of this receptor is too low in mouse ventricular cardiomyocytes, as recently demonstrated,^19^ or the cAMP generated after β_2_-AR activation does not cross the 10-15 nm wide diadic cleft, presumably due to high degree of compartmentalized PDE-mediated cAMP degradation. Subsequent FRET measurements have identified PDE4 as the major PDE type responsible for cAMP hydrolysis in the vicinity of RyR2, while the participation of PDE3 was much less pronounced and that of PDE2 was undetectable in this microdomain (see **Fig. 3**). This is in line with a previous report, which identified PDE4D3 as a direct interaction partner with cAMP hydrolyzing activity in the RyR2 macromolecular complex.^9^

To study the pathological changes of local cAMP dynamics, we focused on a relatively early, often latent cardiac disease state of hypertrophy, because it is significantly less well understood than the chronic and advanced pathology of terminal heart failure. To this end, we used a highly reproducible TAC intervention in the FVB/N1 mouse heart, which has been established as an early disease model of compensated cardiac hypertrophy but without significant reduction of contractile function. From a therapeutic and translational point of view, this compensated stage of latent disease development is especially interesting for testing new pharmacological strategies at the microdomain level. In particular, as previously demonstrated,^4^ this early stage of disease is not yet associated with whole-cell changes of PDE expression and activity (see also **Supplementary Figure 4**). Instead, we were able to uncover a PDE redistribution in or between subcellular microdomains. Previously, we have shown that TAC leads to a redistribution of PDE2 from β_1_-AR to β_2_-AR-associated plasma membrane compartments, while PDE3 activity was specifically reduced at the β_2_-AR with a so far unknown relocation destiny.^4^ In this work, we show that both PDE2 and PDE3 effects are increased at the RyR2 microdomain after TAC (see **Fig. 4**). This suggests that in hypertrophy, PDE3 might redistribute across the cleft from β_2_-AR located at the T-tubular membrane to the junctional SR where RyR2 is found. However, the most prominent disease-associated alteration in the RyR2 microdomain is the dramatic loss of PDE4 activity (see **Figs. 4, 6**), apparently not fully compensated by the increased local PDE2 and PDE3 effects post-TAC. This could be the reason why β_2_-AR/cAMP signals can reach the junctional SR and breach the RyR2 microdomain compartment, similar to healthy cells when PDE4 is inhibited with rolipram. Importantly, in TAC cells or hearts, a local β_2_-AR-induced rise of cAMP at RyR2 could be linked to a significant increase in calcium spark generation and a strong tendency towards increased ventricular arrhythmias (**Fig. 5B, Supplementary Fig. 6**). Importantly, β_2_-AR stimulation triggered calcium waves in TAC but not in sham cells, overall similar to the calcium overload effects associated with β_1_-AR stimulation, which was confirmed in sham cells (see **Fig. 5**). This occurs in concert with a moderate increase of cell contractility and relaxation (see **Supplementary Figure 5**) and might serve as an important pathological mechanism of increased arrhythmia susceptibility in response to normally cardioprotective^25, 26^ β_2_-AR subtype stimulation.

**Figure 6.**
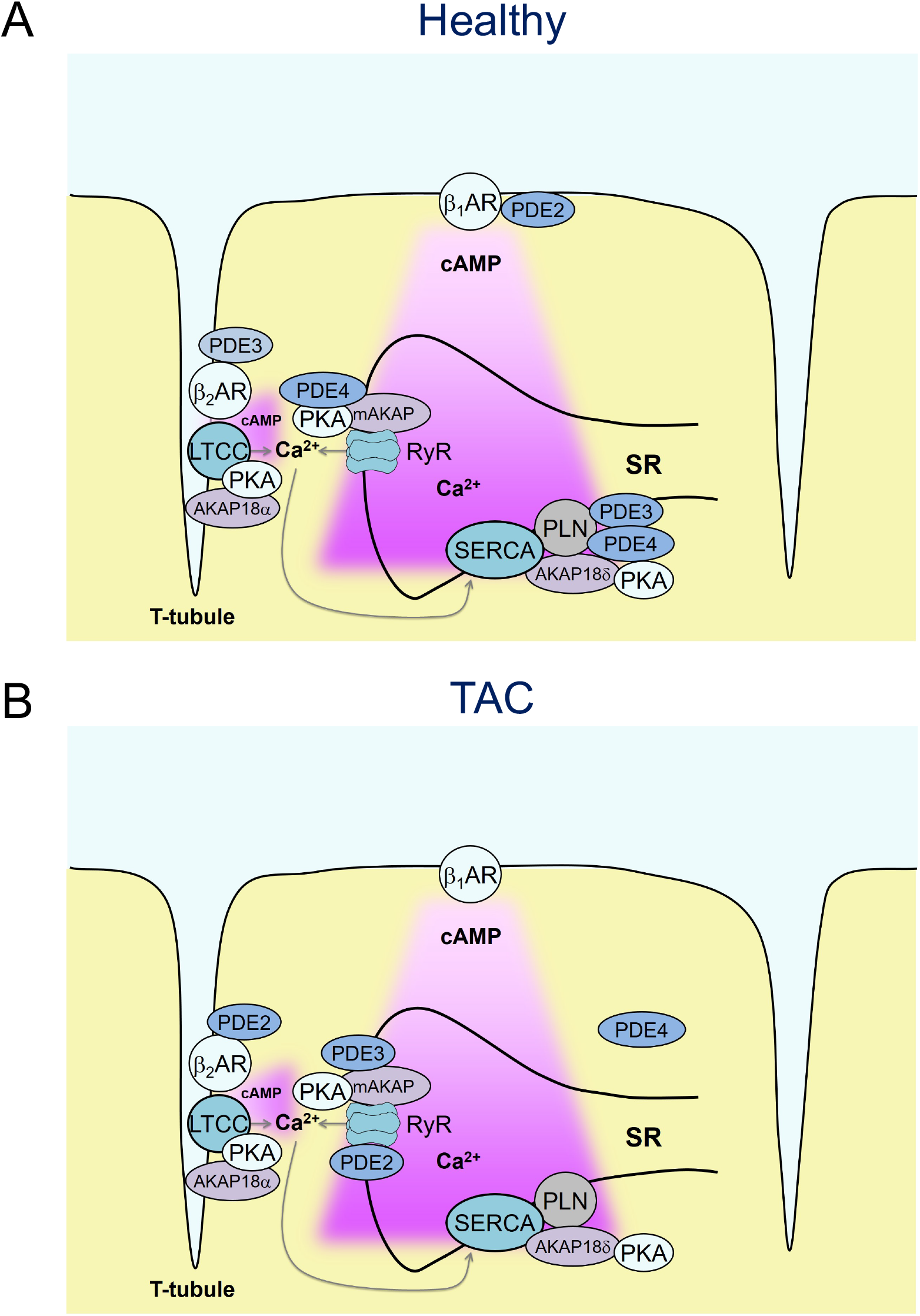
Schematics showing proposed changes in local cAMP signaling occurring in early heart disease. (**A**) In healthy cardiomyocytes, β_1_-but not β_2_-AR-cAMP pools stimulate the RyR2 microdomain because of low β_2_-AR expression and/or tight PDE-mediated control of β_2_-AR/cAMP in the dyadic cleft (**B**) already in early cardiac disease, characterized by hypertrophy but before the onset of chronic heart failure, the RyR2-associated PDE4 effects are lost, which is incompletely compensated by increased local PDE2 and PDE3 contributions. Overall, this might lead to augmented β_2_-AR/cAMP signals at RyR2 channels and increased calcium wave generation in line with increased arrhythmia susceptibility from hypertrophic remodeling.

In conclusion, we have generated a novel FRET-based biosensor and a corresponding transgenic mouse model which can be used to directly monitor local cAMP dynamics in the vicinity of cardiac RyR2. While our experimental data obtained using this system represent only the first step towards understanding local cAMP changes in an early cardiac disease model, this technique clearly demonstrates dramatic alterations of local PDE effects, which might explain significant changes in microdomain specific cAMP levels and increased susceptibility for ventricular arrhythmias. In the future, this real time method can be used to better dissect molecular mechanisms of altered RyR2 function in various cardiac pathologies for the development of new RyR2-targeted therapies aimed at counteracting the detrimental microdomain remodeling.

## Methods

### Chemicals

3-Isobutyl-1-methylxanthin (IBMX) was from AppliChem. BAY 60-7550 cilostamide and rolipram were from Santa Cruz (Santa Cruz, CA). Isoproterenol, forskolin, ICI 118551, CGP 20712A, laminin and all other chemicals were from Sigma-Aldrich (Deisenhofen, Germany).

### Development of the Epac1-JNC biosensor

DNA sequence of the cytosolic cAMP sensor Epac1-camps^27^ without stop codon was fused via a flexible helical linker GSMPLVDFFC to the full-length mouse JNC sequence (excluding the first two amino acids M and A) as shown in Fig. 1a. The resulting construct was subcloned via KpnI and XhoI into a vector containing the α-myosin heavy chain (α-MHC) promoter which was used for generation of transgenic mice.^28^

### Generation of transgenic mice

The α-MHC-based vector containing Epac1-JNC sequence was linearized using SpeI, purified and used for pronuclear injections.^29^ Founder mice and their heterozygote offspring were genotyped by PCR using the following primers: TGACAGACAGATCCCTCCTAT and CATGGCGGACTTGAAGAAGT, giving a ~340 bp fragment on a gel. Positive founder mice were bred with wildtype FVB/NRj animals (Janvier Labs, Saint Berthevin, France) to generate a heterozygous offspring line.

### Cardiomyocyte isolation and FRET measurements

Adult ventricular cardiomyocytes were isolated by enzymatic digestion during Langendorff perfusion, plated on laminin coated round glass coverslides and used for FRET or Ca^2+^ measurements as previously described.^4, 30, 31^

### Transverse aortic constriction (TAC)

All animal experiments were performed in accordance with institutional (Tierschutzbüro Universitätsmedizin Göttingen) and governmental (LAVES Niedersachsen, BGV Hamburg) guidelines. Female mice aged 8-12 weeks were randomized into sham or TAC groups and anesthetized using 2% isoflurane in pure oxygen. A suprasternal incision was made, and the aortic arch was visualized using a binocular operating stereoscope (Olympus). TAC interventions used a spacer-defined (26-gauge) constriction fixed by a 6-0 polyviolene suture between the first and second trunk of the aortic arch.^32^ For sham, the aorta was exposed as for TAC but not constricted. 3 days later, Doppler velocity was measured by a 20 MHz probe to quantify the pressure gradient across the TAC region or after sham procedure by transthoracic echocardiography (VisualSonics Vevo 2100; Toronto, Canada). 3 days before and during 1 week after surgery, mice received analgesic therapy with metamizole in drinking water. 8-10 weeks after surgery, mice were analyzed by echocardiography, hearts were harvested for ventricular cardiomyocyte isolation or histology.^4^

### Immunofluorescence and Confocal Microscopy

wildtype and transgenic Epac1-JNC cells were plated on laminin, fixed with 4 % paraformaldehyde and stained with primary monoclonal RyR2 antibody (MA3-916, Thermo Scientific, 1:500 dilution) followed by the secondary anti-mouse Alexa Fluor 633 (Life Technologies, dilution 1:300). Confocal microscopy was performed using Zeiss LSM710 microscope (Carl Zeiss MicroImaging) equipped with a Plan-Apochromat x63/1.40 oil-immersion objective. Epac1-JNC biosensor fluorescence was excited using the 488 nm diode laser, and Alexa Fluor 633 - using 633 nm diode laser. ZEN 2010 software (Zeiss) was used for image analysis.

### Measurement of Ca^2+^ transients and Ca^2+^ sparks

Ca^2+^ transient measurements were performed as previously described.^33^ Briefly, myocytes were loaded at room temperature with fluo 4-AM (10 μmol/L; Molecular Probes, Eugene, OR) for 30 min, and then another 30 min were allowed for intracellular de-esterification. The composition of the experimental solution was as follows (in mmol/L): 140 NaCl, 5.4 KCl, 0.33 Na_2_HPO_4_, 1.2 MgCl_2_, 10 HEPES, 10 glucose, 1.2 CaCl_2_, pH 7.4 with NaOH. Drugs were added at the indicated concentrations. Confocal microscopy was performed using Zeiss LSM880 microscope (Carl Zeiss MicroImaging) equipped with a Plan-Apochromat x63/1.40 oil-immersion objective. [Ca^2+^]_i_ was measured by fluo-4 epifluorescence with excitation at 488 nm and emission > 500 nm. Line scans were performed at ~2,45 ms time intervals with a 0.1 μm pixels size and applied longitudinally to the cell axis to access the cell length. Data were expressed as ΔF/F0, where F is the fluorescence intensity, and F0 is the intensity at rest. Myocytes were stimulated 5x at 1 Hz via platinum electrodes (5ms, 20V) connected to an IonOptics stimulator (Westwood, MA 02090). Experiments were stopped if the investigated myocyte showed Ca^2+^ waves in control conditions. Ca^2+^ sparks were analyzed after field stimulation by sparkmaster as previously described.^34^

### Statistics

Echocardiographic, morphometric, real time PCR and FRET data were analyzed using the Origin Pro 8.5 software (OriginLab Corporation, Northhampton, MA). Normal distribution was tested by the Kolmogorov-Smirnov test, and differences between the groups were analyzed using mixed ANOVA followed by Wald Chi-squared test, one-way ANOVA for simple two-group comparison or Kruskal-Wallis ANOVA, as appropriate, at the significance level of 0.05. Sample sizes were 3-6 mice and 8-17 measured cells. No sample exclusion was performed, unless in TAC mice which were not studies if the pressure gradients did not reach 50 mmHg (standard cut-off for the development of reliable pressure-overload induced hypertrophy).

## Acknowledgments

We thank the core transgenic unit of the Max-Planck-Institute for Experimental Medicine for transgenic mouse generation, Karina Schlosser for support with histological and immunoblot analysis, Nadja Bork for help with qPCR, and SFB 1002 Service Unit for TAC surgeries and echocardiography.

## References

1. Bers DM. Cardiac excitation-contraction coupling. Nature. 2002;415:198–205.

2. Lompre AM, Hajjar RJ, Harding SE, Kranias EG, Lohse MJ and Marks AR. Ca2+ cycling and new therapeutic approaches for heart failure. Circulation. 2010;121:822–30.

3. Perera RK and Nikolaev VO. Compartmentation of cAMP signalling in cardiomyocytes in health and disease. Acta Physiol (Oxf). 2013;207:650–62.

4. Perera RK, Sprenger JU, Steinbrecher JH, Hubscher D, Lehnart SE, Abesser M, Schuh K, El-Armouche A and Nikolaev VO. Microdomain switch of cGMP-regulated phosphodiesterases leads to ANP-induced augmentation of beta-adrenoceptor-stimulated contractility in early cardiac hypertrophy. Circ Res. 2015;116:1304–11.

5. Sprenger JU, Perera RK, Steinbrecher JH, Lehnart SE, Maier LS, Hasenfuss G and Nikolaev VO. In vivo model with targeted cAMP biosensor reveals changes in receptor-microdomain communication in cardiac disease. Nat Commun. 2015;6:6965.

6. Kokkonen K and Kass DA. Nanodomain Regulation of Cardiac Cyclic Nucleotide Signaling by Phosphodiesterases. Annual Review of Pharmacology and Toxicology, Vol 57. 2017;57:455–479.

7. Froese A and Nikolaev VO. Imaging alterations of cardiomyocyte cAMP microdomains in disease. Front Pharmacol. 2015;6:172.

8. Wehrens XH, Lehnart SE, Reiken SR, Deng SX, Vest JA, Cervantes D, Coromilas J, Landry DW and Marks AR. Protection from cardiac arrhythmia through ryanodine receptor-stabilizing protein calstabin2. Science. 2004;304:292–6.

9. Lehnart SE, Wehrens XH, Reiken S, Warrier S, Belevych AE, Harvey RD, Richter W, Jin SL, Conti M and Marks AR. Phosphodiesterase 4D deficiency in the ryanodine-receptor complex promotes heart failure and arrhythmias. Cell. 2005;123:25–35.

10. Oda T, Yano M, Yamamoto T, Tokuhisa T, Okuda S, Doi M, Ohkusa T, Ikeda Y, Kobayashi S, Ikemoto N and Matsuzaki M. Defective regulation of interdomain interactions within the ryanodine receptor plays a key role in the pathogenesis of heart failure. Circulation. 2005;111:3400–10.

11. Wehrens XH, Lehnart SE, Reiken SR and Marks AR. Ca2+/calmodulin-dependent protein kinase II phosphorylation regulates the cardiac ryanodine receptor. Circ Res. 2004;94:e61–70.

12. Pereira L, Cheng H, Lao DH, Na L, van Oort RJ, Brown JH, Wehrens XH, Chen J and Bers DM. Epac2 mediates cardiac beta1-adrenergic-dependent sarcoplasmic reticulum Ca2+ leak and arrhythmia. Circulation. 2013;127:913–22.

13. Mangmool S, Shukla AK and Rockman HA. beta-Arrestin-dependent activation of Ca(2+)/calmodulin kinase II after beta(1)-adrenergic receptor stimulation. J Cell Biol. 2010;189:573–87.

14. Lehnart SE. Novel targets for treating heart and muscle disease: stabilizing ryanodine receptors and preventing intracellular calcium leak. Curr Opin Pharmacol. 2007;7:225–32.

15. Hong CS, Cho MC, Kwak YG, Song CH, Lee YH, Lim JS, Kwon YK, Chae SW and Kim DH. Cardiac remodeling and atrial fibrillation in transgenic mice overexpressing junctin. FASEB J. 2002;16:1310–2.

16. Kirchhefer U, Neumann J, Bers DM, Buchwalow IB, Fabritz L, Hanske G, Justus I, Riemann B, Schmitz W and Jones LR. Impaired relaxation in transgenic mice overexpressing junctin. Cardiovasc Res. 2003;59:369–79.

17. Bilski AJ, Halliday SE, Fitzgerald JD and Wale JL. The pharmacology of a beta 2-selective adrenoceptor antagonist (ICI 118,551). J Cardiovasc Pharmacol. 1983;5:430–7.

18. Dooley DJ, Bittiger H and Reymann NC. CGP 20712 A: a useful tool for quantitating beta 1- and beta 2-adrenoceptors. Eur J Pharmacol. 1986;130:137–9.

19. Myagmar BE, Flynn JM, Cowley PM, Swigart PM, Montgomery MD, Thai K, Nair D, Gupta R, Deng DX, Hosoda C, Melov S, Baker AJ and Simpson PC. Adrenergic Receptors in Individual Ventricular Myocytes: The Beta-1 and Alpha-1B Are in All Cells, the Alpha-1A Is in a Subpopulation, and the Beta-2 and Beta-3 Are Mostly Absent. Circ Res. 2017;120:1103–1115.

20. Nikolaev VO, Moshkov A, Lyon AR, Miragoli M, Novak P, Paur H, Lohse MJ, Korchev YE, Harding SE and Gorelik J. Beta2-adrenergic receptor redistribution in heart failure changes cAMP compartmentation. Science. 2010;327:1653–7.

21. Calebiro D, Nikolaev VO, Gagliani MC, de Filippis T, Dees C, Tacchetti C, Persani L and Lohse MJ. Persistent cAMP-signals triggered by internalized G-protein-coupled receptors. PLoS Biol. 2009;7:e1000172.

22. Despa S, Shui B, Bossuyt J, Lang D, Kotlikoff MI and Bers DM. Junctional cleft [Ca(2)(+)]i measurements using novel cleft-targeted Ca(2)(+) sensors. Circ Res. 2014;115:339–47.

23. Marx SO, Reiken S, Hisamatsu Y, Jayaraman T, Burkhoff D, Rosemblit N and Marks AR. PKA phosphorylation dissociates FKBP12.6 from the calcium release channel (ryanodine receptor): defective regulation in failing hearts. Cell. 2000;101:365–76.

24. Pritchard TJ and Kranias EG. Junctin and the histidine-rich Ca2+ binding protein: potential roles in heart failure and arrhythmogenesis. J Physiol. 2009;587:3125–33.

25. Communal C, Singh K, Sawyer DB and Colucci WS. Opposing effects of beta(1)- and beta(2)-adrenergic receptors on cardiac myocyte apoptosis : role of a pertussis toxin-sensitive G protein. Circulation. 1999;100:2210–2.

26. Zhu WZ, Zheng M, Koch WJ, Lefkowitz RJ, Kobilka BK and Xiao RP. Dual modulation of cell survival and cell death by beta(2)-adrenergic signaling in adult mouse cardiac myocytes. Proc Natl Acad Sci U S A. 2001;98:1607–12.

27. Nikolaev VO, Bunemann M, Hein L, Hannawacker A and Lohse MJ. Novel single chain cAMP sensors for receptor-induced signal propagation. J Biol Chem. 2004;279:37215–8.

28. Nikolaev VO, Bünemann M, Schmitteckert E, Lohse MJ and Engelhardt S. Cyclic AMP imaging in adult cardiac myocytes reveals far-reaching beta1-adrenergic but locally confined beta2-adrenergic receptor-mediated signaling. Circ Res. 2006;99:1084–91.

29. Buitrago M, Lorenz K, Maass AH, Oberdorf-Maass S, Keller U, Schmitteckert EM, Ivashchenko Y, Lohse MJ and Engelhardt S. The transcriptional repressor Nab1 is a specific regulator of pathological cardiac hypertrophy. Nat Med. 2005;11:837–44.

30. Börner S, Schwede F, Schlipp A, Berisha F, Calebiro D, Lohse MJ and Nikolaev VO. FRET measurements of intracellular cAMP concentrations and cAMP analog permeability in intact cells. Nat Protoc. 2011;6:427–38.

31. Sprenger JU, Perera RK, Götz KR and Nikolaev VO. FRET microscopy for real-time monitoring of signaling events in live cells using unimolecular biosensors. J Vis Exp. 2012:e4081.

32. Hu P, Zhang D, Swenson L, Chakrabarti G, Abel ED and Litwin SE. Minimally invasive aortic banding in mice: effects of altered cardiomyocyte insulin signaling during pressure overload. Am J Physiol Heart Circ Physiol. 2003;285:H1261–9.

33. Brandenburg S, Kohl T, Williams GS, Gusev K, Wagner E, Rog-Zielinska EA, Hebisch E, Dura M, Didie M, Gotthardt M, Nikolaev VO, Hasenfuss G, Kohl P, Ward CW, Lederer WJ and Lehnart SE. Axial tubule junctions control rapid calcium signaling in atria. J Clin Invest. 2016;126:3999–4015.

34. Picht E, Zima AV, Blatter LA and Bers DM. SparkMaster: automated calcium spark analysis with ImageJ. Am J Physiol Cell Physiol. 2007;293:C1073–81.

