## Supplemental Material for "Direct monitoring of cAMP at the cardiac ryanodine receptor using a novel targeted fluorescence biosensor mouse"

#### Supplementary Methods

**Echocardiography, Histology and Morphometric analysis.** Echocardiography of wildtype and transgenic mice was performed at the age of 6 months using the Vevo2100 system (VisualSonics; Toronto, Canada) equipped with a 30 Hz transducer (MS-400 MicroScan Transducer). For histology and morphometric analysis, wildtype and transgenic hearts were harvested, perfused with phosphate-buffered saline until blood free and fixed in 4% Roti Histofix (Roth) at 4°C overnight. Fixed hearts were embedded in paraffin, cut into 5 µm cross sections using Microtom (Leica RM 2165), dewaxed, rehydrated and stained as previously described.[1]

##### Ex-vivo measurement of arrhythmia inducibility

Langendorff-perfused ex-vivo measurements were performed as described previously.[2] Programmed stimulation was applied by using a designated digital stimulus generator (STG4002, Multi Channel Systems, Reutlingen, Germany) at twice the ventricular pacing threshold. Programmed extrastimulus (S1S1: 120 ms, 100 ms, and 80 ms followed by up to 3 extra beats; 60–20 ms with 5 ms stepwise reduction), burst (5 s at S1S1: 50–10 ms with 10 ms stepwise reduction)[3] and miniburst pacing protocols (S1-S1: 20x 100ms, S2: 3-10 from 60 to 20ms with 2ms decrements)[4] was used to detect arrhythmia susceptibility. Counted were ventricular tachycardias (VTs) defined as ≥4 consecutive premature ventricular complexes.

#### Supplementary References


1. Perera RK, Sprenger JU, Steinbrecher JH, Hubscher D, Lehnart SE, Abesser M, Schuh K, El-Armouche A and Nikolaev VO. Microdomain switch of cGMP-regulated phosphodiesterases leads to ANP-induced augmentation of beta-adrenoceptor-stimulated contractility in early cardiac hypertrophy. Circ Res. 2015;116:1304-11.

2. Jungen C, Scherschel K, Eickholt C, Kuklik P, Klatt N, Bork N, Salzbrunn T, Alken F, Angendohr S, Klene C, Mester J, Klocker N, Veldkamp MW, Schumacher U, Willems S, Nikolaev VO and Meyer C. Disruption of cardiac cholinergic neurons enhances susceptibility to ventricular arrhythmias. Nat Commun. 2017;8:14155.

3. Jungen C, Scherschel K, Bork NI, Kuklik P, Eickholt C, Kniep H, Klatt N, Willems S, Nikolaev VO and Meyer C. Impact of Intracardiac Neurons on Cardiac Electrophysiology and Arrhythmogenesis in an Ex Vivo Langendorff System. J Vis Exp. 2018; 22:135.

4. Clasen L, Eickholt C, Angendohr S, Jungen C, Shin DI, Donner B, Furnkranz A, Kelm M, Klocker N, Meyer C and Makimoto H. A modified approach for programmed electrical stimulation in mice: Inducibility of ventricular arrhythmias. PLoS One. 2018;13:e0201910.

5. Börner S, Schwede F, Schlipp A, Berisha F, Calebiro D, Lohse MJ and Nikolaev VO. FRET measurements of intracellular cAMP concentrations and cAMP analog permeability in intact cells. Nat Protoc. 2011;6:427-38.

#### Supplementary Tables

**Supplementary Table 1. Echocardiographic phenotyping of the wildtype vs. Epac1-JNC transgenic mice at 6 months of age**

| Parameter | Genotype | |
|---|---|---|
| | **Wildtype** | **Transgenic** |
| ATWThD (mm) | 0.80 ± 0.01 | 0.86 ± 0.02* |
| ATWThS (mm) | 1.23 ± 0.04 | 1.25 ± 0.05 |
| PWThD (mm) | 0.80 ± 0.01 | 0.85 ± 0.02* |
| PWThS (mm) | 1.22 ± 0.04 | 1.24 ± 0.05 |
| LV-EDD (mm) | 4.5 ± 0.1 | 4.4 ± 0.1 |
| LV-ESD (mm) | 3.3 ± 0.1 | 3.3 ± 0.1 |
| FS (%) | 26.2 ± 1.7 | 26.4 ± 1.8 |
| FAS (%) | 43.9 ± 2.1 | 42.9 ± 2.4 |
| EF (%) | 49.8 ± 2.2 | 48.0 ± 2.2 |
| HR (bpm) | 458 ± 20 | 471 ± 14 |
| n | 12 | 10 |

Shown are means ± SE. AWTh, anterior wall thickness in diastole (D) and systole (S); PWTh, posterior wall thickness in diastole (D) and systole (S); LV-EDD, left ventricular end-diastolic dimension; LV-ESD, left ventricular end-systolic dimension; FS, fractional shortening; FAS, fractional area shortening; EF, ejection fraction; HR, heart rate; bpm, beats per minute; n, number of mice analyzed per group. * $p<0.05$ by one-way ANOVA. All other parameters were not significantly different between the two groups at $p=0.05$ by one-way ANOVA.

**Supplementary Table 2. Echocardiographic parameters for Epac1-JNC mice 8 weeks after sham or TAC surgery**

| Parameter | Surgery | |
|---|---|---|
| | Sham | TAC |
| Gradient (mmHg) | 4.8 ± 0.5 | 66.4± 4.3* |
| ATWThD (mm) | 0.89 ± 0.07 | 1.22 ± 0.08* |
| ATWThS (mm) | 1.32 ± 0.07 | 1.67 ± 0.06* |
| PWThD (mm) | 0.78 ± 0.04 | 1.00 ± 0.03* |
| PWThS (mm) | 1.15 ± 0.04 | 1.35 ± 0.07* |
| LV-EDD (mm) | 4.1 ± 0.1 | 3.8 ± 0.1 |
| LV-ESD (mm) | 2.8 ± 0.1 | 2.6 ± 0.1 |
| FS (%) | 32.4 ± 2.6 | 33.1 ± 2.6 |
| FAS (%) | 50.5 ± 3.3 | 51.3 ± 2.7 |
| EF (%) | 56.5 ± 2.8 | 58.1 ± 2.5 |
| HR (bpm) | 467 ± 17 | 445 ± 17 |
| HW/BW | 4.1 ± 0.3 | 5.4 ± 0.3 |
| n | 10 | 12 |

Shown are means ± SE. AWTh, anterior wall thickness in diastole (D) and systole (S); PWTh, posterior wall thickness in diastole (D) and systole (S); LV-EDD, left ventricular end-diastolic dimension; LV-ESD, left ventricular end-systolic dimension; FS, fractional shortening; FAS, fractional area shortening; EF, ejection fraction; HR, heart rate; bpm, beats per minute; HW/BW, calculated heart weight to body weight ratio; n, number of mice analyzed per group. * $p<0.05$ by one-way ANOVA. All other parameters were not significantly different between the two groups at $p=0.05$ by one-way ANOVA.

**Supplementary Table 3. Echocardiographic parameters for FVB/N1 mice 8 weeks after sham or TAC surgery**

| Parameter | Surgery | |
|---|---|---|
| | Sham | TAC |
| Gradient (mmHg) | 4.9 ± 1.1 | 81.7 ± 4.9* |
| ATWThD (mm) | 1.12 ± 0.05 | 1.31 ± 0.09* |
| ATWThS (mm) | 1.59 ± 0.04 | 1.69 ± 0.07 |
| PWThD (mm) | 0.78 ± 0.02 | 1.03 ± 0.07* |
| PWThS (mm) | 1.08 ± 0.01 | 1.35 ± 0.11* |
| LV-EDD (mm) | 4.1 ± 0.1 | 4.1 ± 0.1 |
| LV-ESD (mm) | 2.7 ± 0.1 | 2.9 ± 0.1 |
| FS (%) | 33.4 ± 1.1 | 30.0 ± 1.4 |
| FAS (%) | 55.0 ± 2.2 | 47.5 ± 2.6 |
| EF (%) | 62.6 ± 2.7 | 51.7 ± 2.4* |
| HR (bpm) | 462 ± 33 | 489 ± 11 |
| HW/BW | 4.2 ± 0.1 | 5.0 ± 0.3* |
| n | 4 | 4 |

Shown are means ± SE. AWTh, anterior wall thickness in diastole (D) and systole (S); PWTh, posterior wall thickness in diastole (D) and systole (S); LV-EDD, left ventricular end-diastolic dimension; LV-ESD, left ventricular end-systolic dimension; FS, fractional shortening; FAS, fractional area shortening; EF, ejection fraction; HR, heart rate; bpm, beats per minute; HW/BW, calculated heart weight to body weight ratio; n, number of mice analyzed per group. * $p<0.05$ by one-way ANOVA. All other parameters were not significantly different between the two groups at $p=0.05$ by one-way ANOVA.

### Supplementary Figure 1 (Berisha et al.)

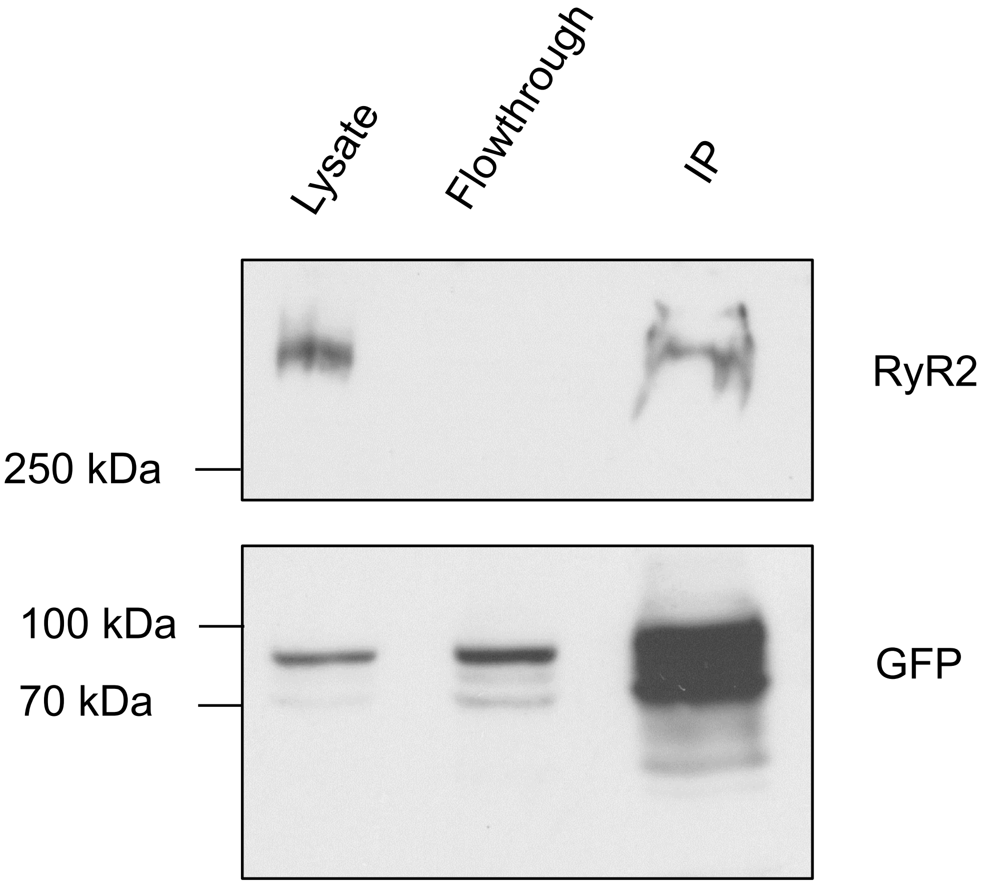


**Supplementary Figure 1. Co-immunoprecipitation of the Epac1-JNC biosensor with cardiac ryanodine receptor.** Epac1-JNC was pulled down from transgenic heart lysates using GFP-Trap® and immunoprecipitated samples were probed for RyR2 and GFP. Representative experiment (n=3).

### Supplementary Figure 2 (Berisha et al.)

A

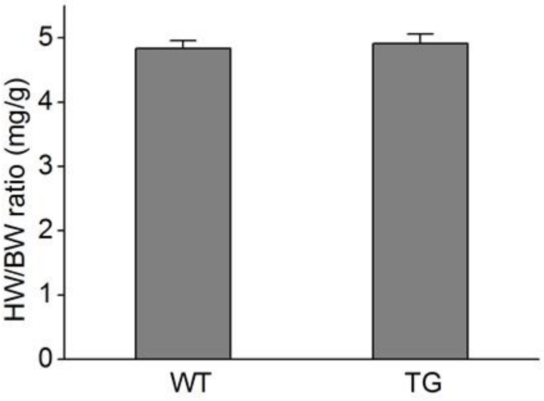


B

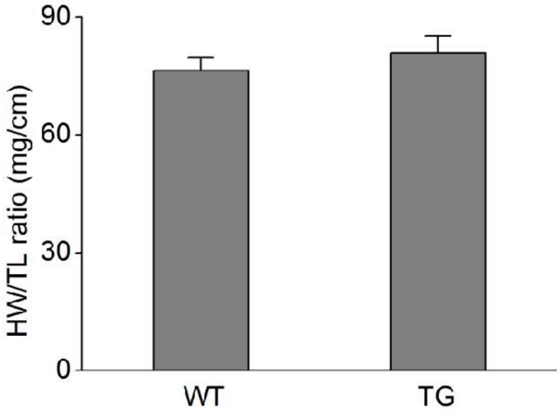


C

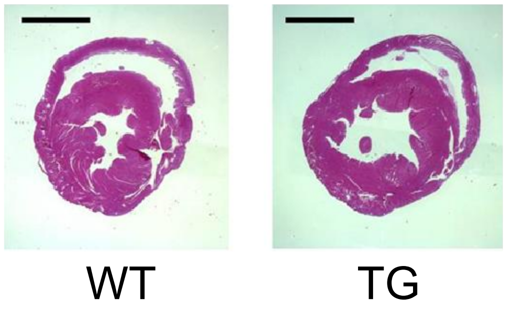


D

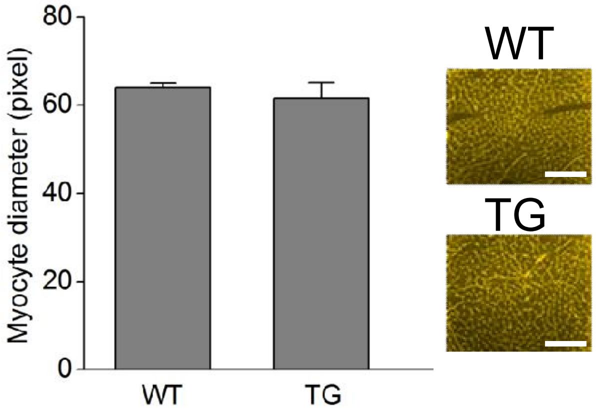


E

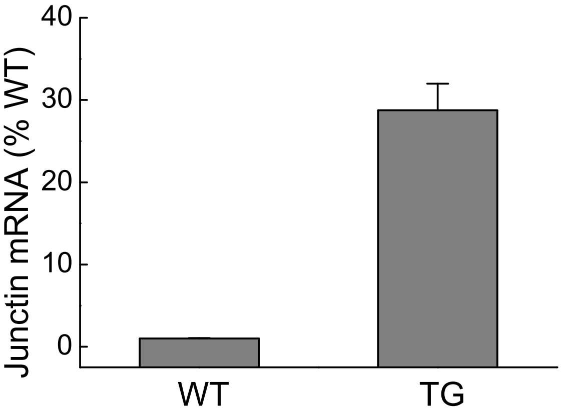


**Supplementary Figure 2. Analysis of morphometric parameters, myocyte size and junctin over-expression. (A)** Heart-weight-to-body-weight (HW/BW) and heart-weight-to-tibia-length (HW/TL) ratios **(B)** at the age of 6 months (means ± SE, n= 13-14 each) show no differences between wildtype (WT) and transgenic (TG) animals. **(C)** Representative (n>5) hematoxylin and eosin stained heart cross sections at the age of 6 months. Scale bars, 2 mm. **(D)** Diameter of transgenic (TG) Epac1-JNC cardiomyocytes was not different compared to wildtype (WT) cells at the age of 6 months as investigated by WGA assay. Data are means ± SE from 6 WT and 4 TG hearts with 100 cells counted per each section. Scale bars, 100 µm. **(E)** qPCR analysis of junctin mRNA expression in TG cardiomyocytes as compared to endogenous wildtype junctin. Means ± SE (n=5 mice each).

### Supplementary Figure 3 (Berisha et al.)

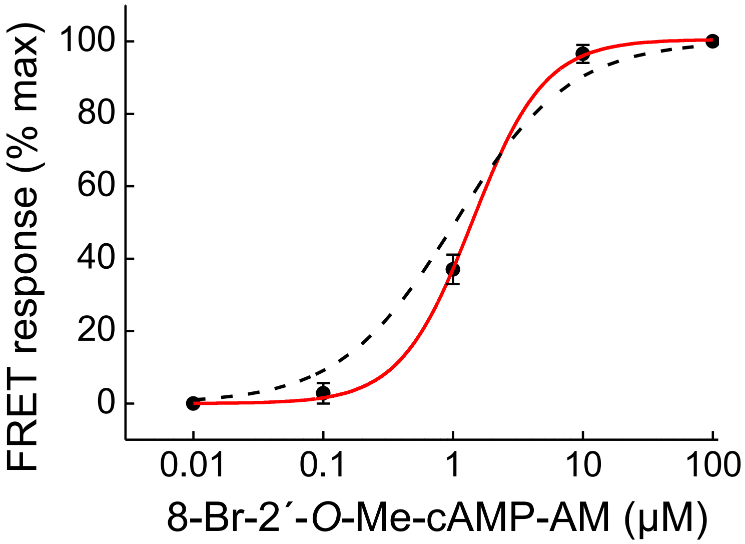


**Supplementary Figure 3. Concentration-response dependency of Epac1-JNC sensor in myocytes.** Transgenic cardiomyocytes were pretreated with the adenylyl cyclase inhibitor MDL12,330A (100 µM) for 10 minutes and stimulated with increasing concentrations of the cell-permeable cAMP analogue cAMP 8-Br-2'-O-Me-cAMP-AM. This analogue can be used for direct in cell sensor calibration since it has the same affinity for the sensor binding domains as native cAMP (see Supplementary Ref. 5). Previously published concentration-response dependency for the parental cytosolic sensor Epac1-camps (Supplementary Ref. 1) is shown as dotted line. Epac1-JNC has a comparable cAMP affinity as the cytosolic Epac1-camps sensor. $EC_{50}$ values were 1.2±0.5 and 1.2±0.1 µM (n=4, means ± SE), respectively.

### Supplementary Figure 4 (Berisha et al.)

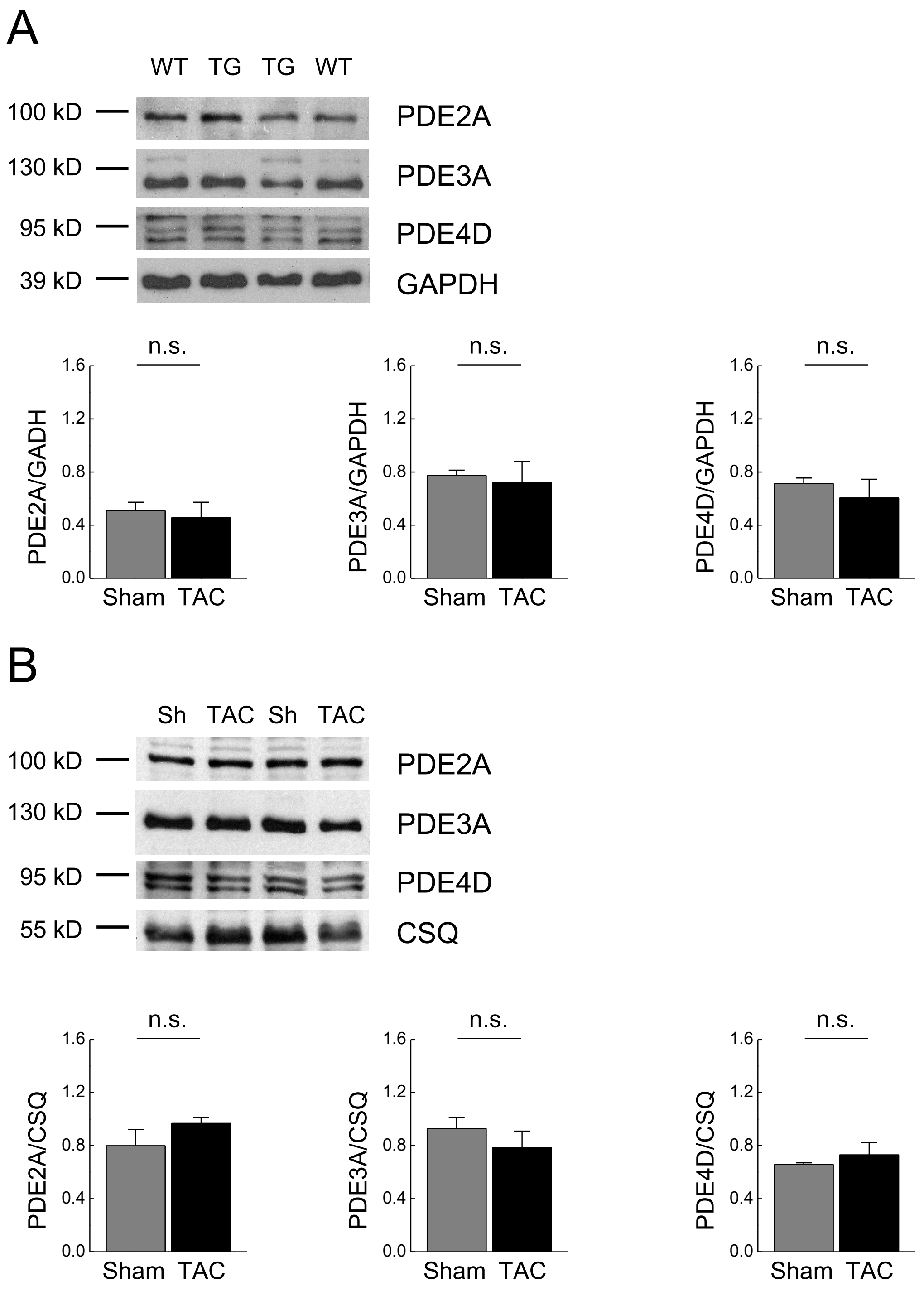


**Supplementary Figure 4. Immunoblot analysis of major PDE families in wildtype and Epac1-JNC myocytes. (A)** Cardiomyocytes were isolated from wildtype (n=4) and transgenic (n=4) mice, lysed and probed with PDE2A, PDE3A and PDE4D antibodies. Representative blots and data analysis for PDE expression shown as PDE/GAPDH, means ± SE. **(B)** Cardiomyocytes were isolated from Epac1-JNC mice 8 weeks after sham or TAC surgery and probed for the same PDEs. Representative blots and data analysis (means ± SE, n=3 each).

### Supplementary Figure 5 (Berisha et al.)

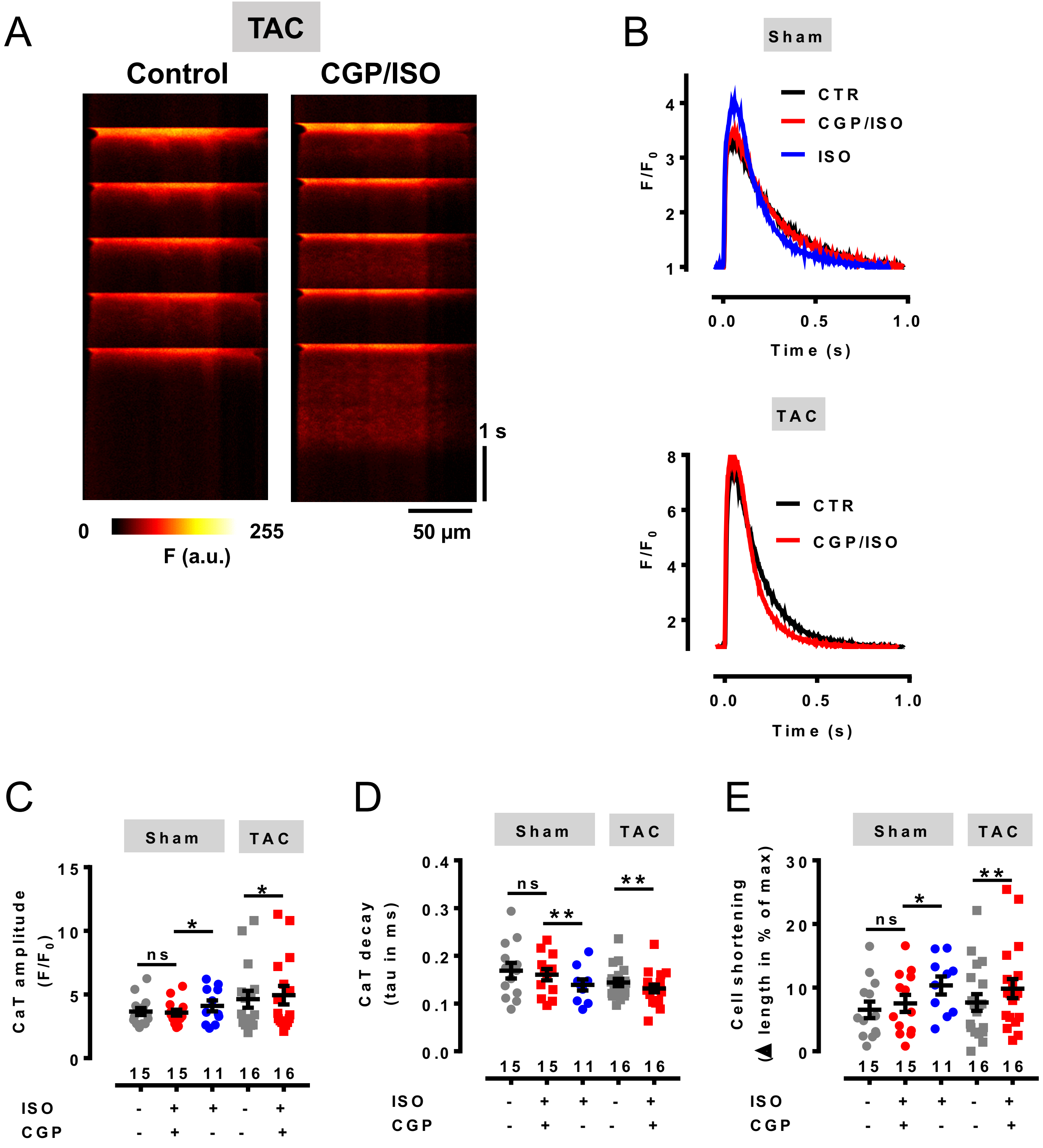


**Supplementary Figure 5. Analysis of electrically evoked $Ca^{2+}$ transients in cardiomyocytes from sham and TAC mice. (A)** Confocal line-scan images in a ventricular cardiomyocyte from a TAC- mouse under unstimulated control conditions (left) and in the presence of ISO plus CGP20712A (right) stimulated at 1 Hz. **(B)** Typical records of spatially averaged $Ca^{2+}$ transients in a myocyte from a sham (left) and a TAC (right) mouse. $Ca^{2+}$ transients recorded in control conditions, in the presence of ISO alone, or in the presence of ISO plus CGP20712A are superimposed. **(C)** Amplitudes of $Ca^{2+}$ transients in myocytes from sham and TAC mice. **(D)** Decay of $Ca^{2+}$ transients in myocytes from sham and TAC mice. The decay was fitted by an exponential function from peak to baseline and expressed as the time constant tau. **(E)** Cell shortening of myocytes isolated from sham and TAC mice induced by electrical stimulation at 1 Hz. Cell shortening was calculated from the length of the fluorescent signal during field stimulation from the confocal line-scan images. Drugs indicated were used at 100 nM each. Statistically significant differences were calculated by paired Student's t-test and indicated with *, $p<0.05$; **, $p<0.01$; ***, $p<0.001$; ns, non-significant. Crosses indicate means ± SEM. Numbers correspond to the number of cells isolated from 3 mice per group.

### Supplementary Figure 6 (Berisha et al.)

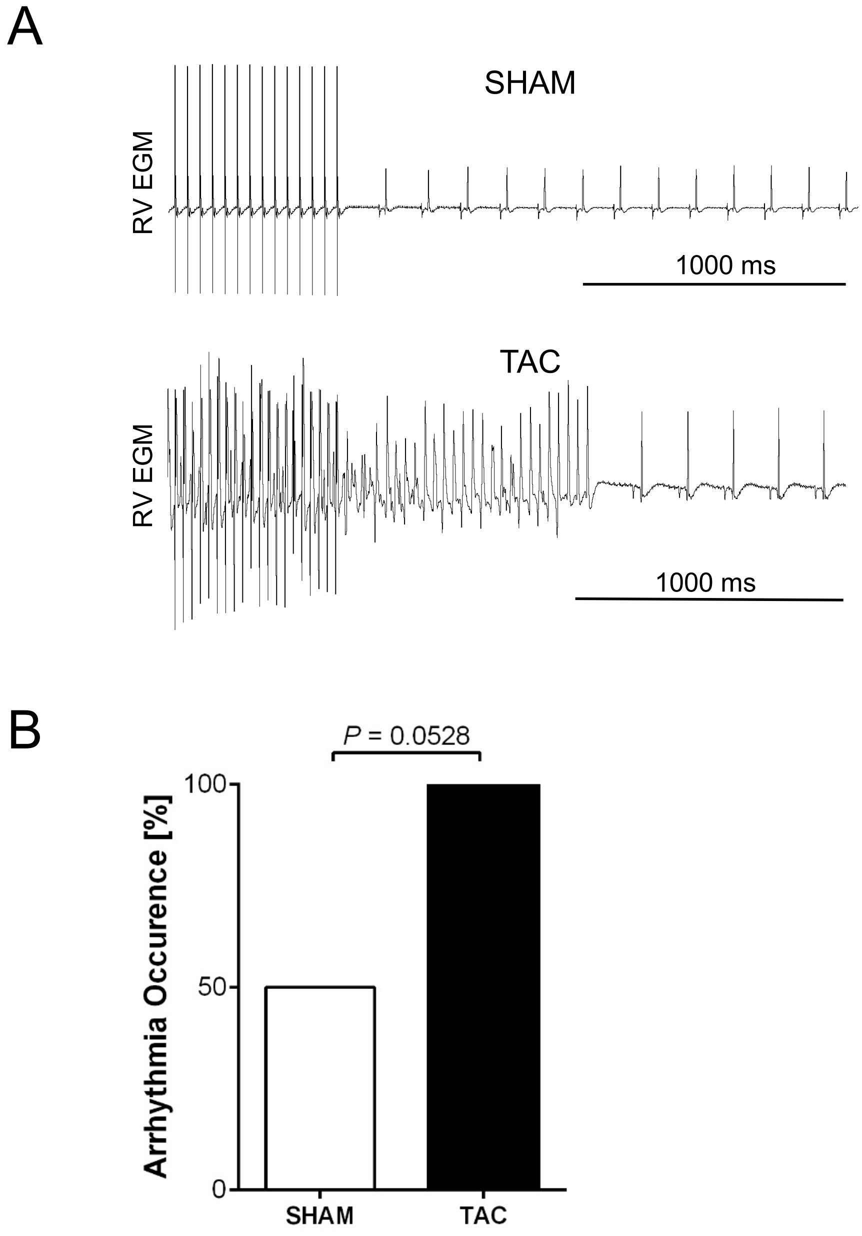


**Supplementary Figure 6. Measurements of arrhythmia occurrence in Langendorff perfused hearts. (A)** Hearts were equilibrated for 15-20 min, pharmacologically stimulated for 10 min with 100 nmol/L ISO in presence of 100 nmol/L CGP20712A, and subjected to a complex stimulation protocol described in Supplementary Methods. Shown are exemplary right ventricular electrograms (RV EGM) tracings from sham and TAC hearts. **(B)** Arrhythmia occurrence was measured as number of hearts with VTs. TAC hearts showed elevated arrhythmia susceptibility. Data are from 4 sham and 6 TAC hearts, $\chi^2$ test.
